## Supplemental Materials for "Epidemiological modeling of SARS-CoV-2 in white-tailed deer (*Odocoileus virginianus*) reveals conditions for introduction and widespread transmission"

Running title: SARS-CoV-2 outbreaks in white-tailed deer

PROPOSED AUTHOR LIST: ELIAS ROSENBLATT[1,*], JONATHAN D. COOK[2], GRAZIELLA V. DIRENZO[3,4], EVAN H.C. GRANT[5], FERNANDO ARCE[4], KIM M. PEPIN[6], F. JAVIERA RUDOLPH[2,7], MICHAEL C. RUNGE[2], SUSAN SHRINER[6], DANIEL WALSH[8], BRITTANY A. MOSHER[1]

[1] *Rubenstein School of Environment and Natural Resources, University of Vermont, Burlington, VT, USA*

[2] *U.S. Geological Survey, Eastern Ecological Science Center, Laurel, MD, USA*

[3] *U. S. Geological Survey, Massachusetts Cooperative Fish and Wildlife Research Unit, University of Massachusetts, Amherst, MA, USA*

[4] *Department of Environmental Conservation, University of Massachusetts, Amherst, MA, USA*

[5] *U.S. Geological Survey, Eastern Ecological Science Center, Turner's Falls, MA, USA*

[6] *National Wildlife Research Center, USDA, APHIS, Fort Collins, CO, USA*

[7] *Department of Ecosystem Sciences and Management, Pennsylvania State University, University Park, PA, USA*

[8] *U. S. Geological Survey, Montana Cooperative Wildlife Research Unit, University of Montana, Missoula, MT, USA*

Supporting Tables and Figures

**Table S1**: Names and affiliations of experts involved in expert elicitation for a study of epidemiological modeling of SARS-CoV-2 in white-tailed deer (*Odocoileus virginianus*) The panel they served on is indicated.

| Expert Name | Affiliation | Expert Panel |
|---|---|---|
| Kamen Campbell | Mississippi Department of Wildlife, Fisheries, and Parks | Deer Ecology |
| Dr. Jeffery Chandler | USDA APHIS National Wildlife Research Center | Virology |
| Dr. Paul Cross | USGS Northern Rocky Mountain Science Center | Virology |
| Dr. Anna Fagre | Centers for Disease Control | Virology |
| Dr. Sarah Hamer | Texas A&M University | Virology |
| Dr. Kate Huyvaert | Washington State University | Virology |
| Dr. Christopher Jennelle | Minnesota Department of Natural Resources | Deer Ecology |
| Dr. Jeff Root | USDA APHIS National Wildlife Research Center | Virology |
| Noelle Thompson | Kentucky Department of Fish and Wildlife Resources | Deer Ecology |
| Jonathan Trudeau | Maryland Department of Natural Resources | Deer Ecology |
| Dr. Kurt Vandegrift | Pennsylvania State University | Deer Ecology |

**Supporting Materials S1: Expert elicitation methods**

From September – December 2022, these two expert panels responded to two sets of questions specific to their discipline and each met for two group discussions. The first set of questions were training questions focused on parameters already estimated in the literature, with the intent of familiarizing experts with the elicitation Shiny platform (Chang et al. 2023) and the four-point elicitation process and building expert confidence in their ability as an individual and group to capture a likely range of parameter values.

After answering and discussing the training questions, experts answered questions on the parameters of interest. After submitting initial estimates, each expert panel met to facilitate knowledge sharing and discuss clarifications to reduce linguistic uncertainty. Estimates from the four-point elicitation process were fit to a log-normal or logit-normal distribution depending on the nature of each parameter. Following these discussions, experts were then invited to submit revised estimates.

Using finalized parameter estimates for each expert, we fit a log-normal or logit-normal distribution to each response and then averaged across common quantiles in R (Howerton et al. 2023; R Core Team 2023). Lastly, we fit an aggregate distribution using the *qmedist*() function from the fitdistrplus package (Delignette-Muller and Dutang 2015).

Any use of trade, firm, or product names is for descriptive purposes only and does not imply endorsement by the U.S. Government.

**Supplemental Text S2: Descriptions of settings for Deer Ecology panel expert elicitation**
Three scenarios were described to experts serving on the Deer Ecology expert panel to contextualize the questions presented to them. This contextualization was essential for reducing linguistic uncertainty within and between experts.

*Setting 1: Wild deer in a rural setting (Questions 4-8, informing Figures S4-S8)*

Consider a wild deer population with the following characteristics to answer the following question. This free-ranging population has a density of 7.7 deer/km$^2$ (20 deer/mi$^2$). During fall months (September-December) a deer will have an average of 17 proximity events with other wild deer each day. The habitat is typical of the agro-forested midwestern region of the United States, comprised of 40-54% agriculture and 46-60% forested.

Assume that human density in this scenario is approximately 3.1 humans/km$^2$, or 8 humans/mi$^2$ (typical median density for a rural midwestern U.S. area). Assume that regulated hunting occurs and that hunters use typical harvest methods, including still- and stand/blind based hunting. Baiting and backyard feeding is illegal but may still occur. We define a “proximity event” as an instance in which two deer come within 1.5 m of each other (or a human and a deer come within 1.5 m of each other.) We define “direct contact” as a condition in which two deer make direct physical contact, including mucous membrane contact (through licking, grooming, or mating).

*Setting 2: Wild deer in a suburban setting (Questions 9-10, informing Figures S9-S10)*

Now consider a wild population of deer in a suburban setting. This free-ranging population has a density of 7.7 deer/km$^2$ (20 deer/mi$^2$). Assume this population shares the landscape with a human population of 100 humans/km$^2$, or 259 humans/mi$^2$.

We define a "proximity event" as an instance in which two deer come within 1.5 m of each other (or a human and a deer come within 1.5 m of each other.) We define "direct contact" as a condition in which two deer make direct physical contact, including mucous membrane contact (through licking, grooming, or mating).

*Setting 3: Captive deer in a intensive facility setting (Questions 11-13, informing Figures S11-S13)*

Consider a captive deer facility used for captive breeding, meat production, or exhibition, with supplemental feeding and periodic corralling and herding of deer to move between areas for facility operations. These herds are stocked at higher densities relative to wild herds or captive herds kept for hunting opportunities. Human interactions with deer range from the movement of deer between paddocks or facilities, direct handling of deer for breeding purposes and veterinary care, and direct handling by visitors.

We define a "proximity event" as an instance in which two deer come within 1.5 m of each other (or a human and a deer come within 1.5 m of each other.)

**Figure S1**: Responsesby experts on the Virology panel to Question 1 to estimate immunity loss rate (α) — Consider a healthy individual white-tailed deer that was recently infected with SARS-CoV-2 and has since recovered (i.e., the individual is no longer shedding infectious virions). Assume that, with recovery, this individual is temporarily immune to reinfection by SARS-CoV-2 if they were to be exposed to the virus by a dose otherwise sufficient to cause infection. However, after some period, this individual will lose their immunity and become susceptible to infection again. <u>After how many days can this individual deer be reinfected with SARS-CoV-2 after it has fully recovered from an infection?</u> (A) fitted log-normal probability distributions for answers provided by individual experts, and (B) the aggregated log-normal distribution of answers across experts (black line). The aggregate log-normal distribution has a median of 112.6 days (80% confidence interval: 50.5 – 251.4 days; grey range along x-axis).

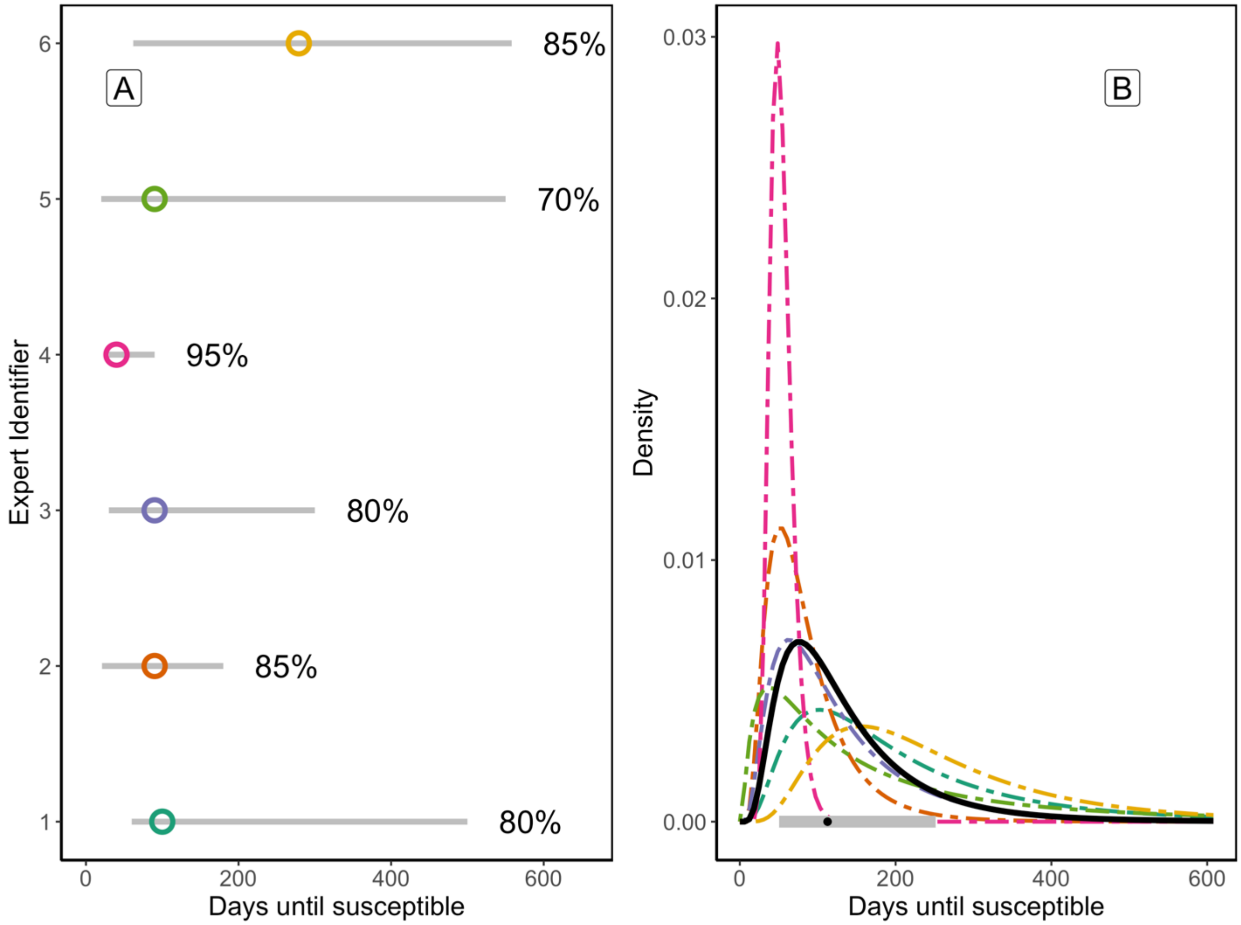

**Figure S2**: Responses by experts on the Virology panel to Question 2 to estimate viral load in deer sputum ($C_{v\text{-deer}}$) — Consider a white-tailed deer infected with SARS-CoV-2 from which you can collect a sputum sample. What is the ratio of the average viral load of a deer compared to the average viral load in an infected human's sputum sample? (A) fitted log-normal probability distributions for answers from individual experts, and (B) the aggregated log-normal distribution of answers across experts. The aggregate log-normal distribution has a median viral load of 1.24 that found in humans (80% confidence interval: 0.80 – 1.93; grey range along x-axis). The vertical lines in both (A) and (B) refer to a ratio of 1, that corresponds to no difference between viral loads in deer and human sputum.

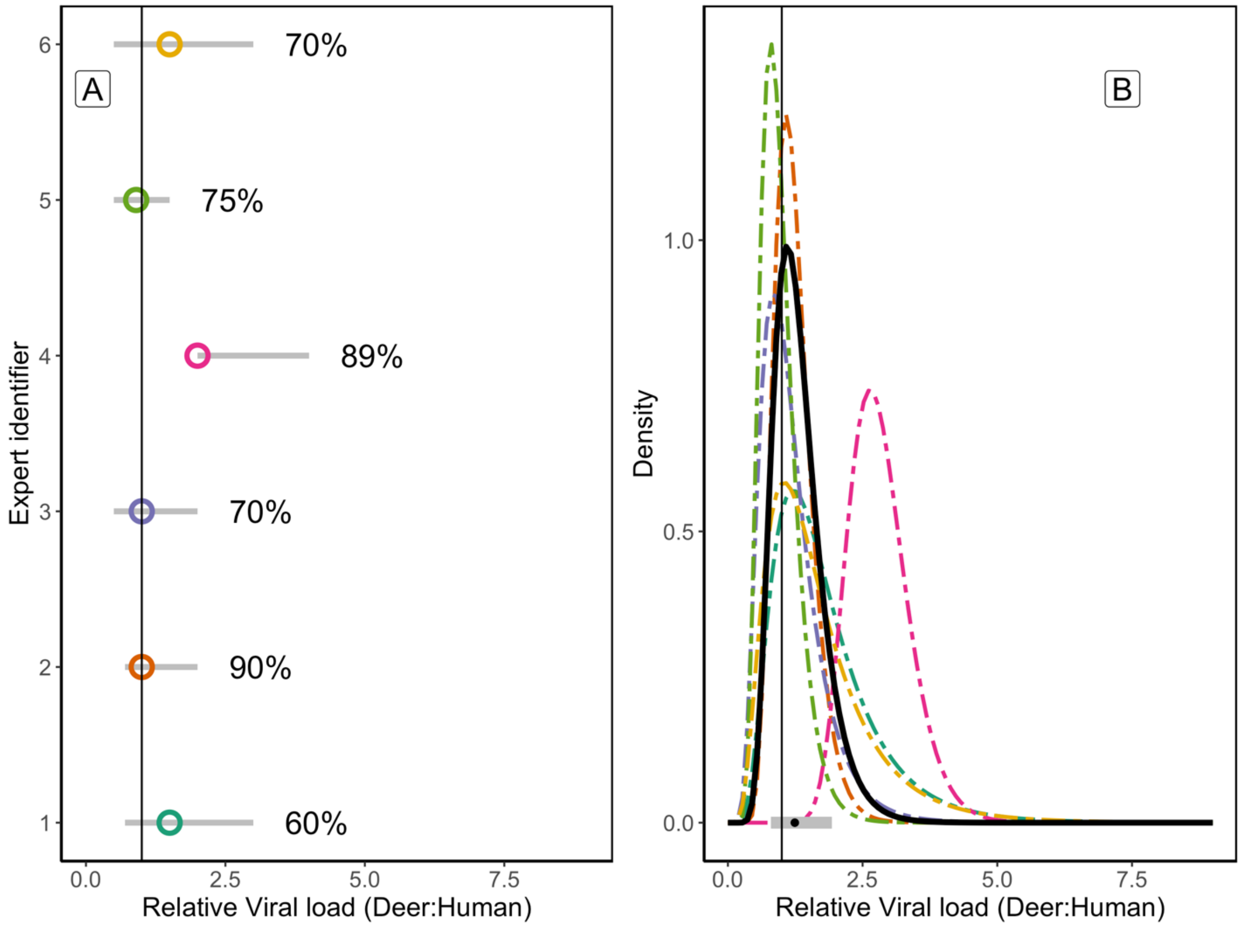

**Figure S3**: Responses by experts on the Virology panel to Question 3 to estimate aerosolized dose-response relationship for deer and SARS-CoV-2 ($\theta$) — For humans, $\theta$=1, corresponding to a 1 quantum dose successfully infecting 63% of susceptible individuals, or HID63. Based on your expertise and knowledge of the literature, what do you expect the r value to be for the average, healthy white-tailed deer? (A) fitted log-normal probability distributions for answers from individual experts, and (B) the aggregated log-normal distribution of answers across experts. The aggregate log-normal distribution has a median dose-relationship of 1.32 (80% confidence interval: 0.93 – 1.87; grey range along x-axis). The vertical line indicates the human dose-response relationship, $\theta$ = 1.

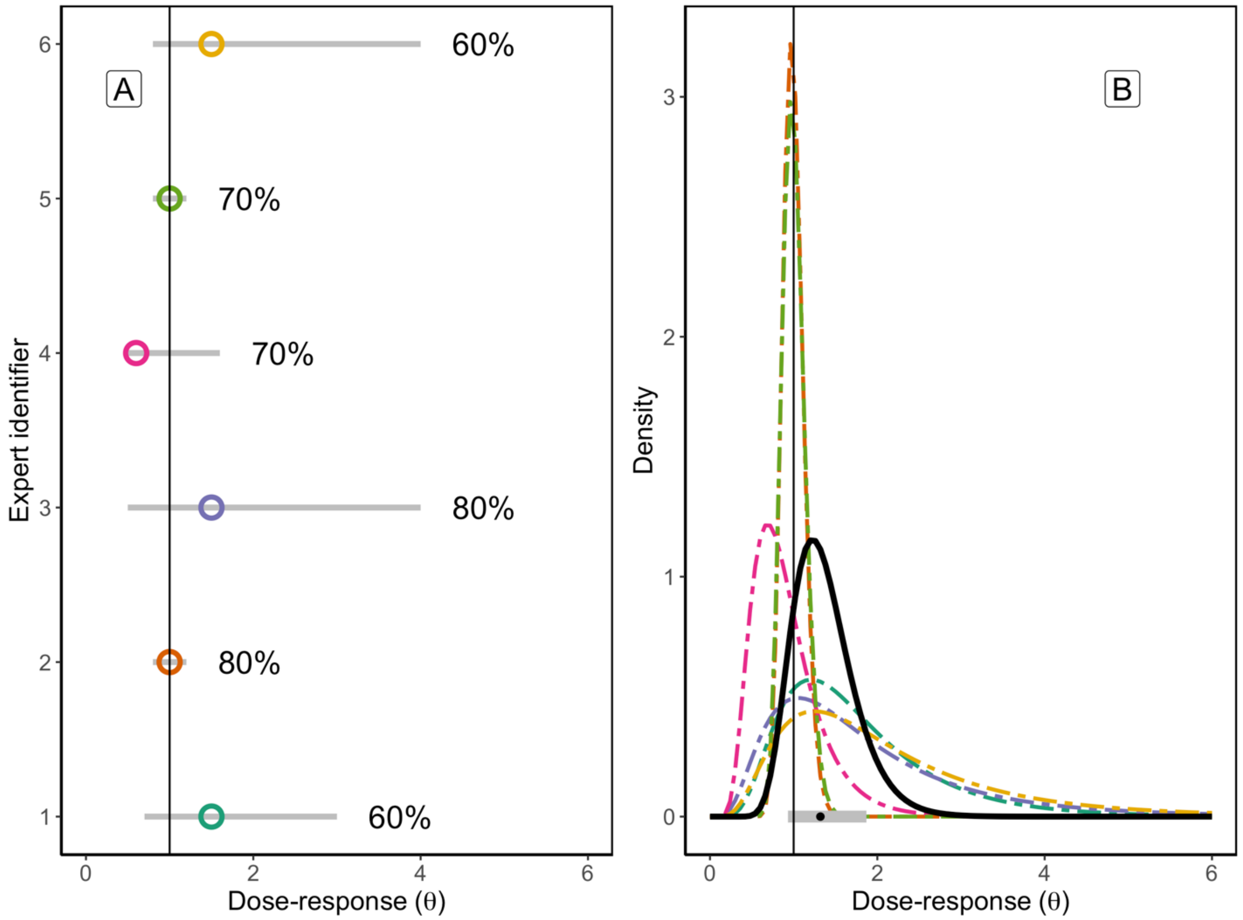

**Figure S4**: Responses by experts on the Deer Ecology panel to Question 4 to estimate the duration of deer staying in proximity of each other (<1.5m; $t_{contact}$) — Given that two individual deer are in proximity (within 1.5 m of each other), how long do you expect these individuals to stay in proximity on average (minutes)? (A) fitted log-normal probability distributions for answers from individual experts, and (B) the aggregated log-normal distribution of answers across experts. The aggregate log-normal distribution has a median duration of 4.72 minutes (80% confidence interval: 0.93 – 24.11 minutes; grey range along x-axis).

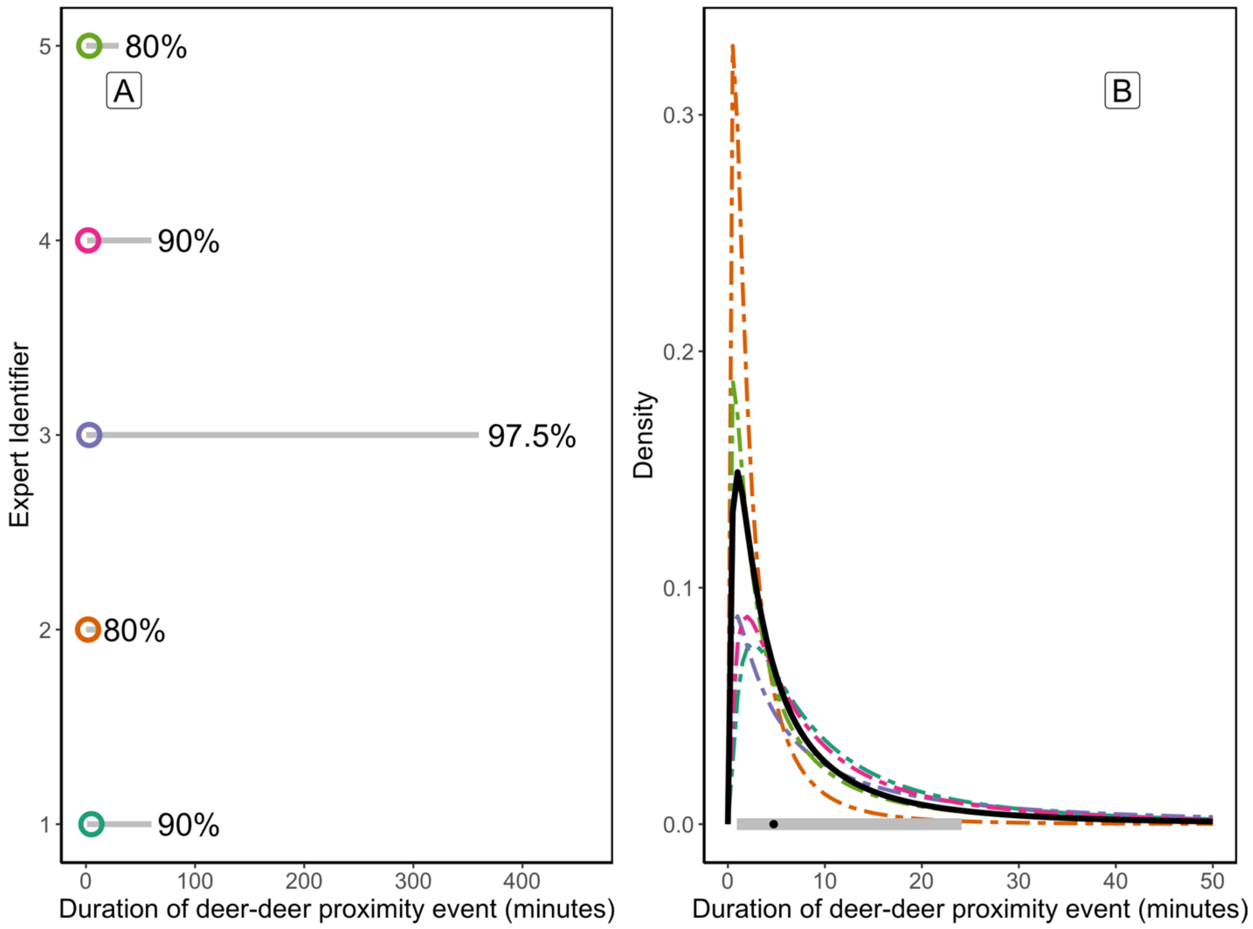

**Figure S5**: Responses by experts on the Deer Ecology panel to Question 5 to estimate the probability of direct contact given proximity ($\varepsilon^{DC}$) — Given that two deer are in proximity (within 1.5m of each other), what is the probability that these individuals make direct contact? (A) fitted logit-normal probability distributions for answers from individual experts, and (B) the aggregated logit-normal distribution of answers across experts. The aggregate logit-normal distribution has a median direct contact probability of 0.19 (80% confidence interval: 0.09 – 0.37; grey range along x-axis).

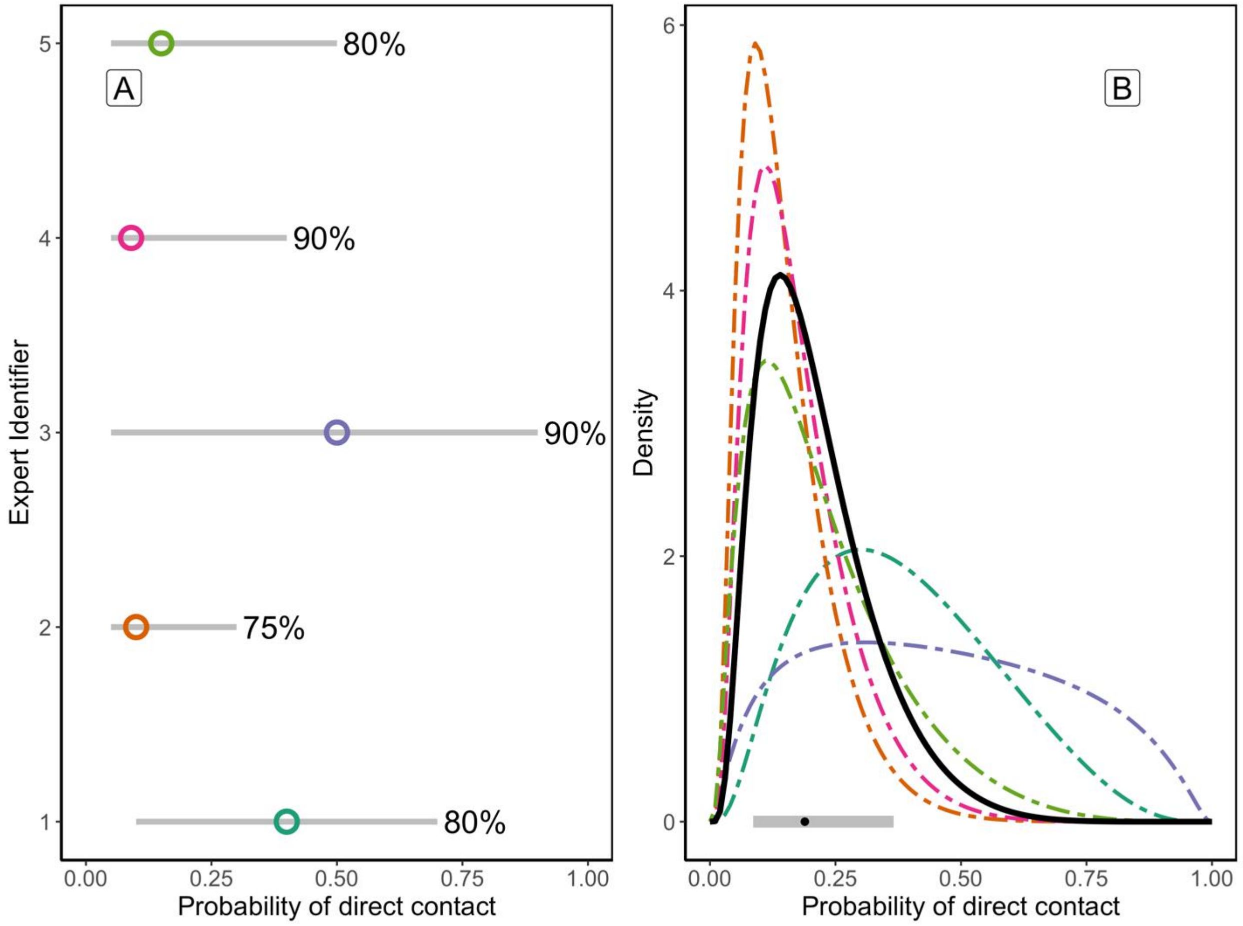

**Figure S6**: Responses by experts on the Deer Ecology panel to Question 6 to estimate the influence of baiting or supplemental feeding ($\rho_{attractant}$) — If an individual deer has 17 proximity events with other deer each day in the absence of baiting, how many proximity events do you expect an individual deer to have with other wild deer in the presence of an attractant (bait, food, or other product intended to attract deer)? (A) fitted log-normal probability distributions for answers from individual experts, and (B) the aggregated log-normal distribution of answers across experts. The aggregate log-normal distribution has a median 32.2 proximity events per day when an attractant is present (80% confidence interval: 24.1 – 43.0). Relative to 17 proximity events in the absence of baiting, this aggregate distribution estimates an increase in proximity events by 1.90-fold when an attractant is present (80% confidence interval: 1.42 – 2.53; grey range along x-axis).

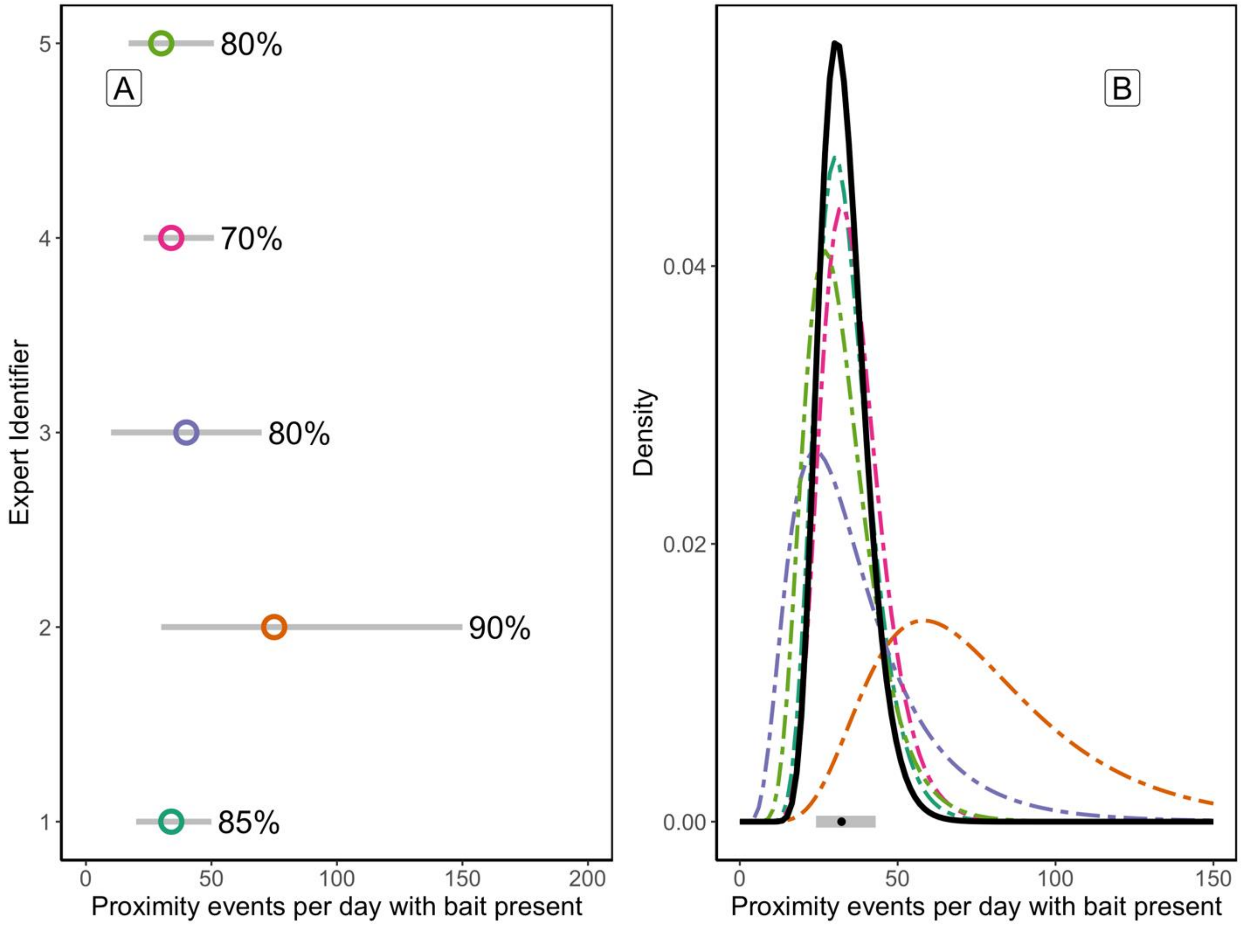

**Figure S7**: Responses by experts on the Deer Ecology panel to Question 7 to estimate the rate of a deer in proximity to a human in a rural setting ($\omega_{HW-rural}$) Given the conditions outlined above [Supplemental Text S1], how many times do you expect an individual deer to come into proximity with a human during the fall months (1 September – 31 December)? (A) fitted log-normal probability distributions for answers from individual experts, and (B) the aggregated log-normal distribution of answers across experts. The aggregate log-normal distribution has a median of 0.20 proximity events per deer, per fall in a rural setting (80% confidence interval: 0.02–1.80; grey range along x-axis).

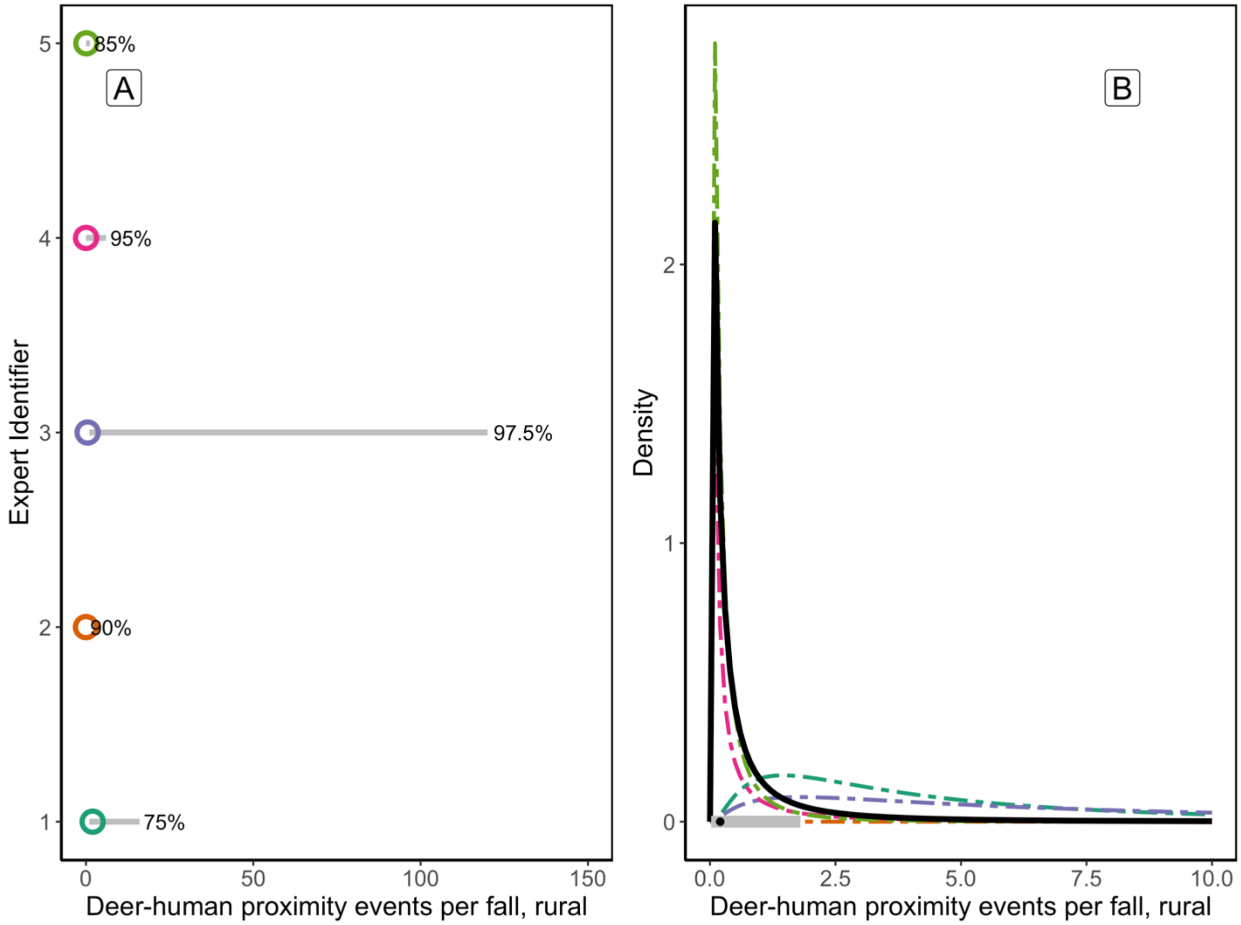

**Figure S8**: Responses by experts on the Deer Ecology panel to Question 8 to estimate the duration of deer staying in proximity of a human in a rural setting (<1.5m; $t_{contact\text{-}HW,\ rural}$). Given that a human and a deer come into proximity in a rural setting (within 1.5m of each other), how long do you expect these individuals (human and deer) to stay in proximity on average (minutes)? (A) fitted log-normal probability distributions for answers from individual experts, and (B) the aggregated log-normal distribution of answers across experts. The aggregate log-normal distribution has a median proximity duration of 0.70 minutes in a rural setting (80% confidence interval: 0.20–2.46 minutes; grey range along x-axis).

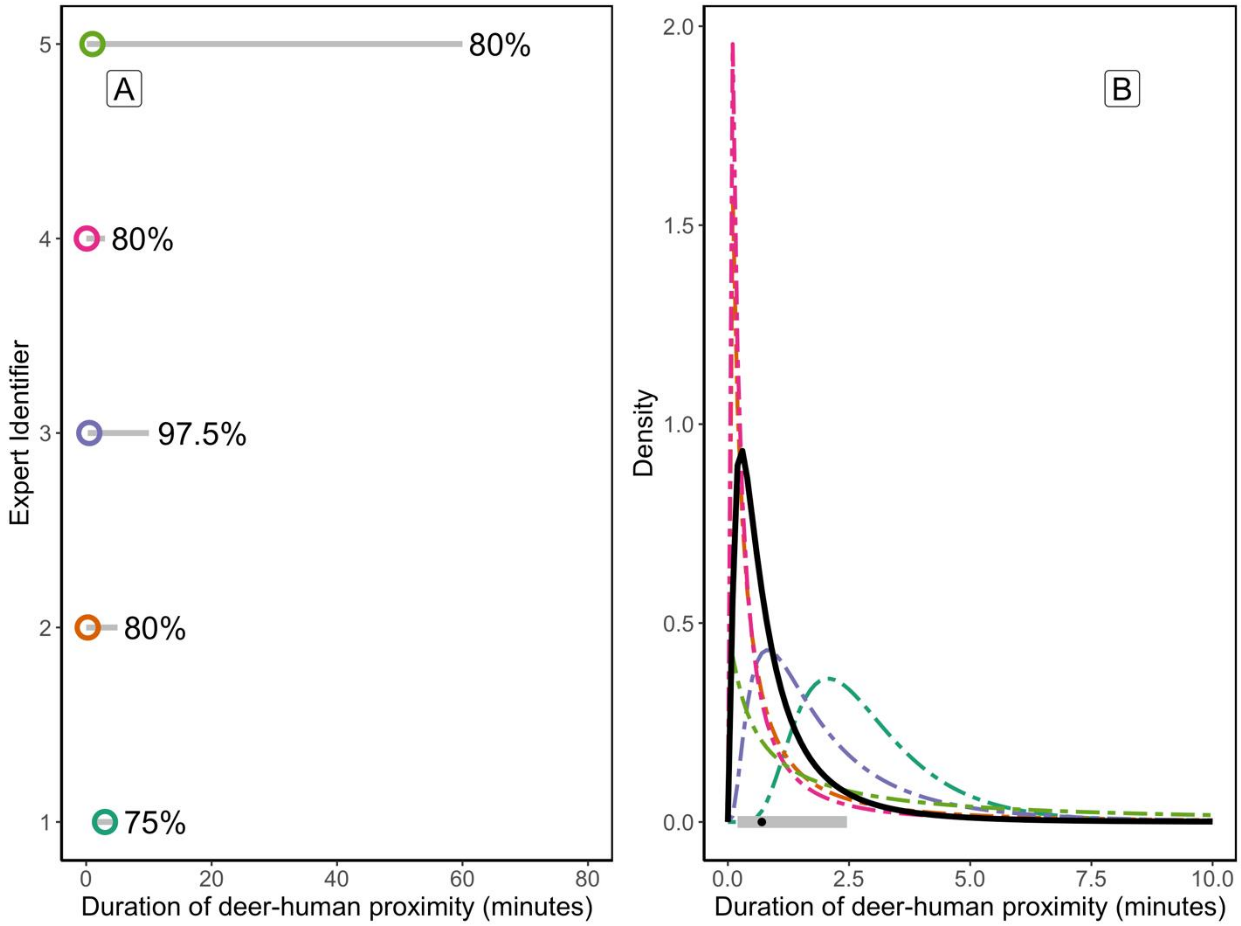

**Figure S9**: Responses by experts on the Deer Ecology panel to Question 9 to estimate the rate of a deer in proximity to a human in a suburban setting ($\omega_{HW-suburban}$) Given the conditions outlined above [Supplemental Text S1], how many times do you expect an individual deer to come into proximity with a human during the fall months (1 September – 31 December)? (A) fitted log-normal probability distributions for answers from individual experts, and (B) the aggregated log-normal distribution of answers across experts. The aggregate log-normal distribution has a median of 1.77 proximity events per deer, per fall in a suburban setting (80% confidence interval: 0.52–6.00; grey range along x-axis).

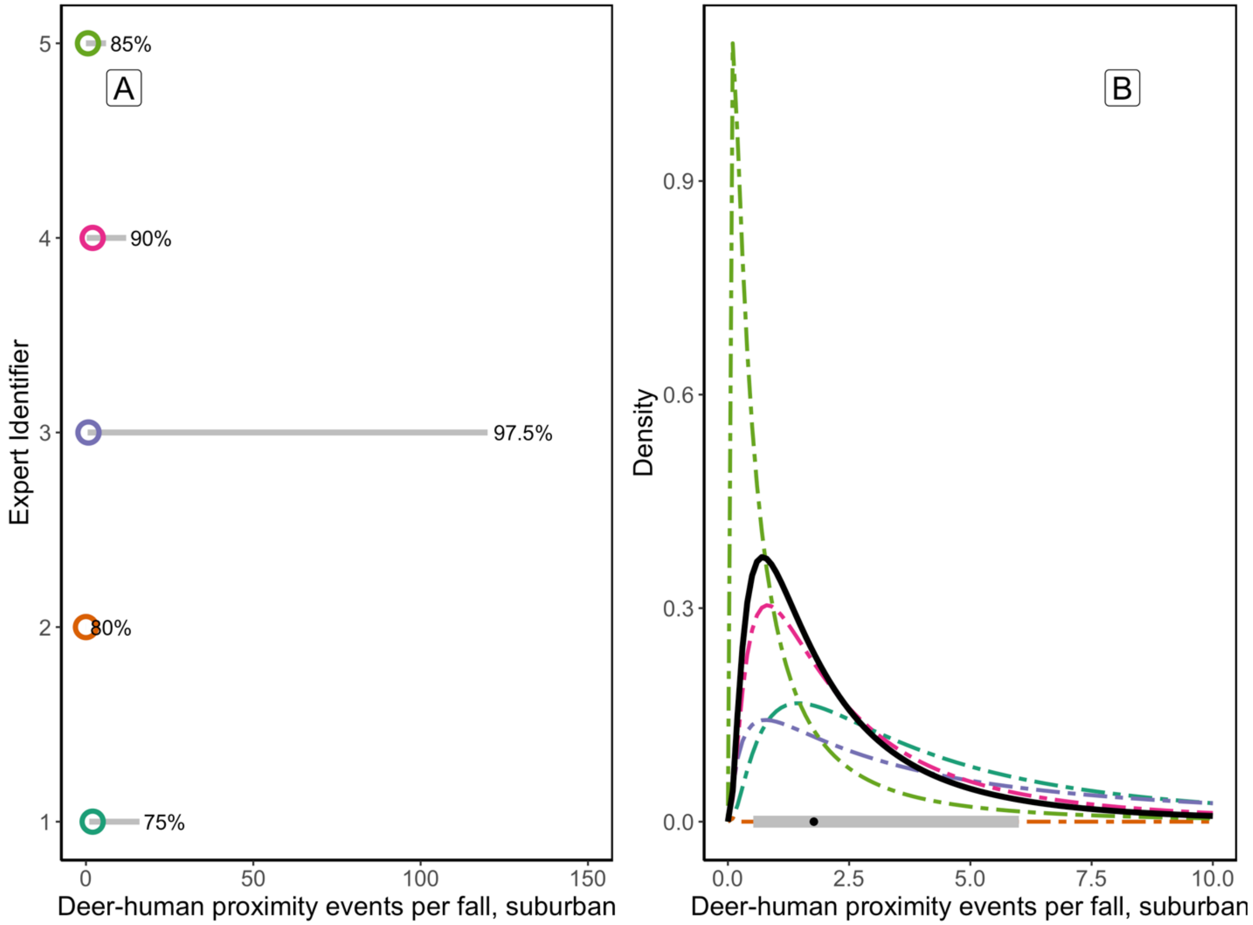

**Figure S10**: Responses by experts on the Deer Ecology panel to Question 10 to estimate the duration of deer staying in proximity of a human in a suburban setting (<1.5m; $t_{contact\text{-}HW,\ suburban}$). <u>Given that a human and a deer come into proximity in a suburban setting (within 1.5m of each other), how long do you expect these individuals (human and deer) to stay in proximity on average (minutes)?</u> (A) fitted log-normal probability distributions for answers from individual experts, and (B) the aggregated log-normal distribution of answers across experts. The aggregate log-normal distribution has a median proximity duration of 1.54 minutes in a suburban setting (80% confidence interval: 0.47–5.07 minutes; grey range along x-axis).

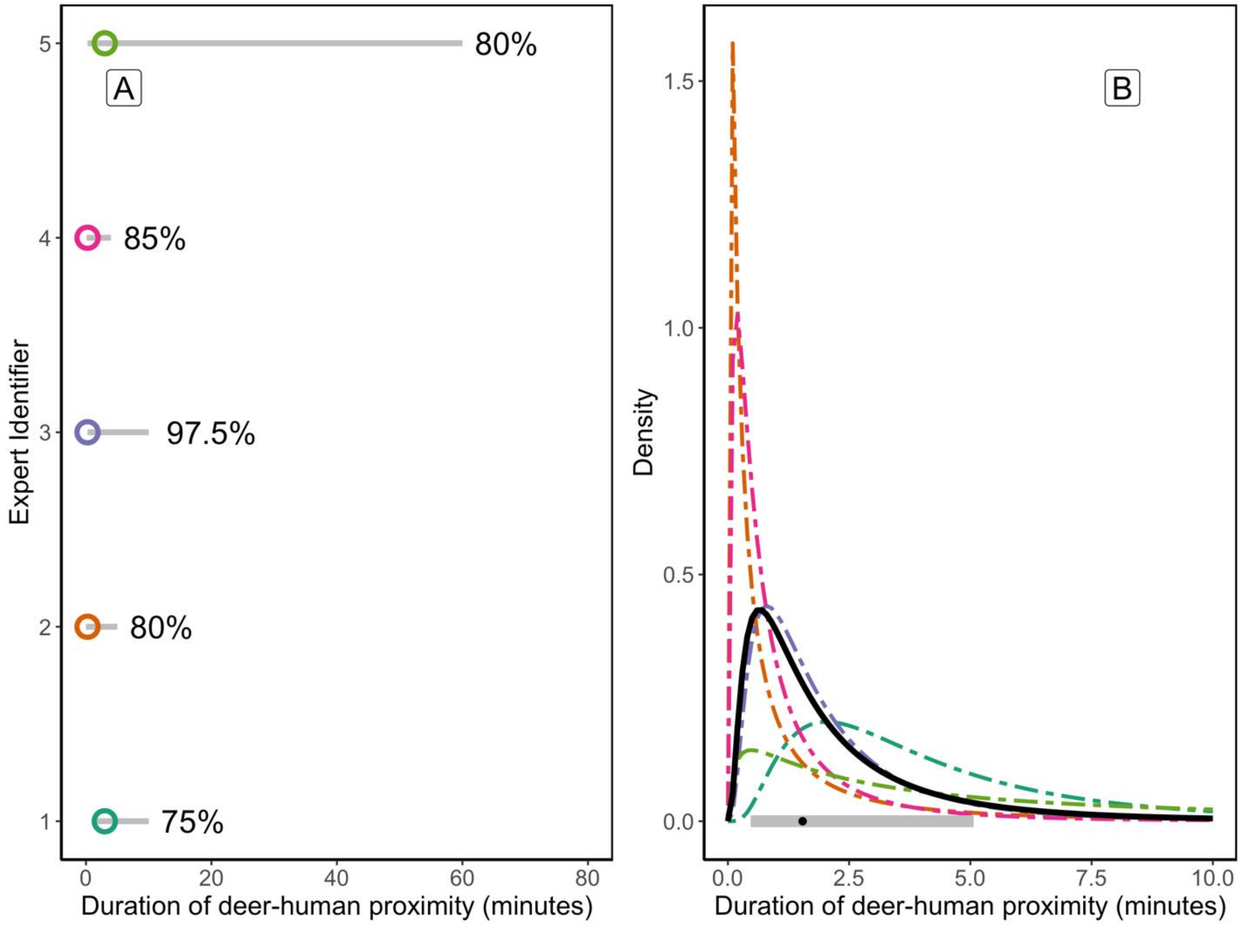

**Figure S11**: Responses by experts on the Deer Ecology panel to Question 11 to estimate the rate of a deer in proximity to a human in an intensive captive setting ($\omega_{HC}$). <u>How many times do you expect an individual deer in a captive facility to come into proximity with a human (within 1.5m of each other) during the fall months (1 September – 31 December)?</u> (A) fitted log-normal probability distributions for answers from individual experts, and (B) the aggregated log-normal distribution of answers across experts. The aggregate log-normal distribution has a median of 12.44 proximity events per deer, per fall in an intensive captive setting (80% confidence interval: 2.92–53.08; grey range along x-axis).

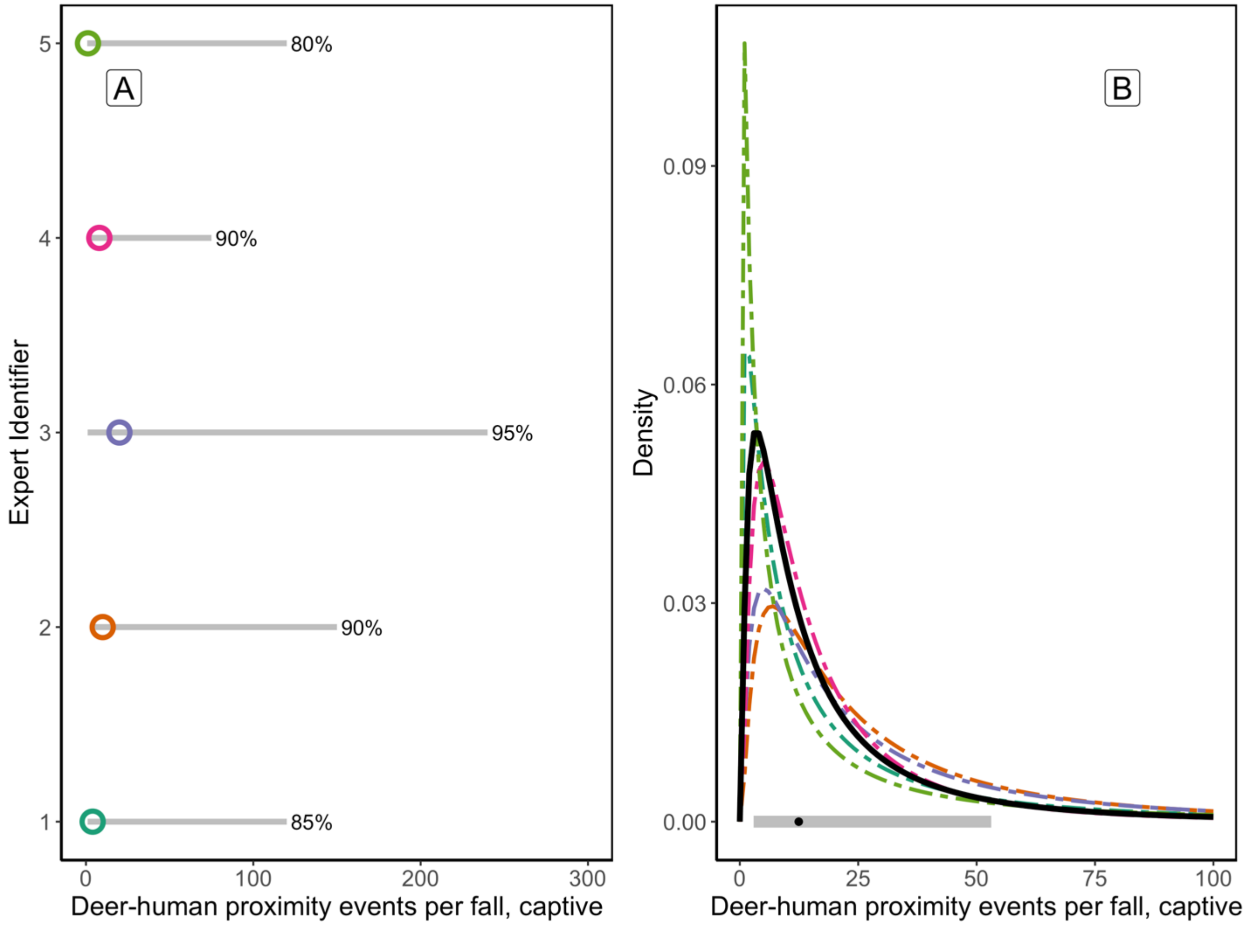

**Figure S12**: Responses by experts on the Deer Ecology panel to Question 12 to estimate the duration of deer staying in proximity of a human in a intensive captive setting (<1.5m; $t_{contact\text{-}CW}$). <u>Given that a human and a deer come into proximity in a captive setting (within 1.5m of each other), how long do you expect these individuals (human and deer) to stay in proximity on average (minutes)?</u> (A) fitted log-normal probability distributions for answers from individual experts, and (B) the aggregated log-normal distribution of answers across experts. The aggregate log-normal distribution has a median proximity duration of 5.98 minutes in an intensive captive setting (80% confidence interval: 1.36–26.16 minutes; grey range along x-axis).

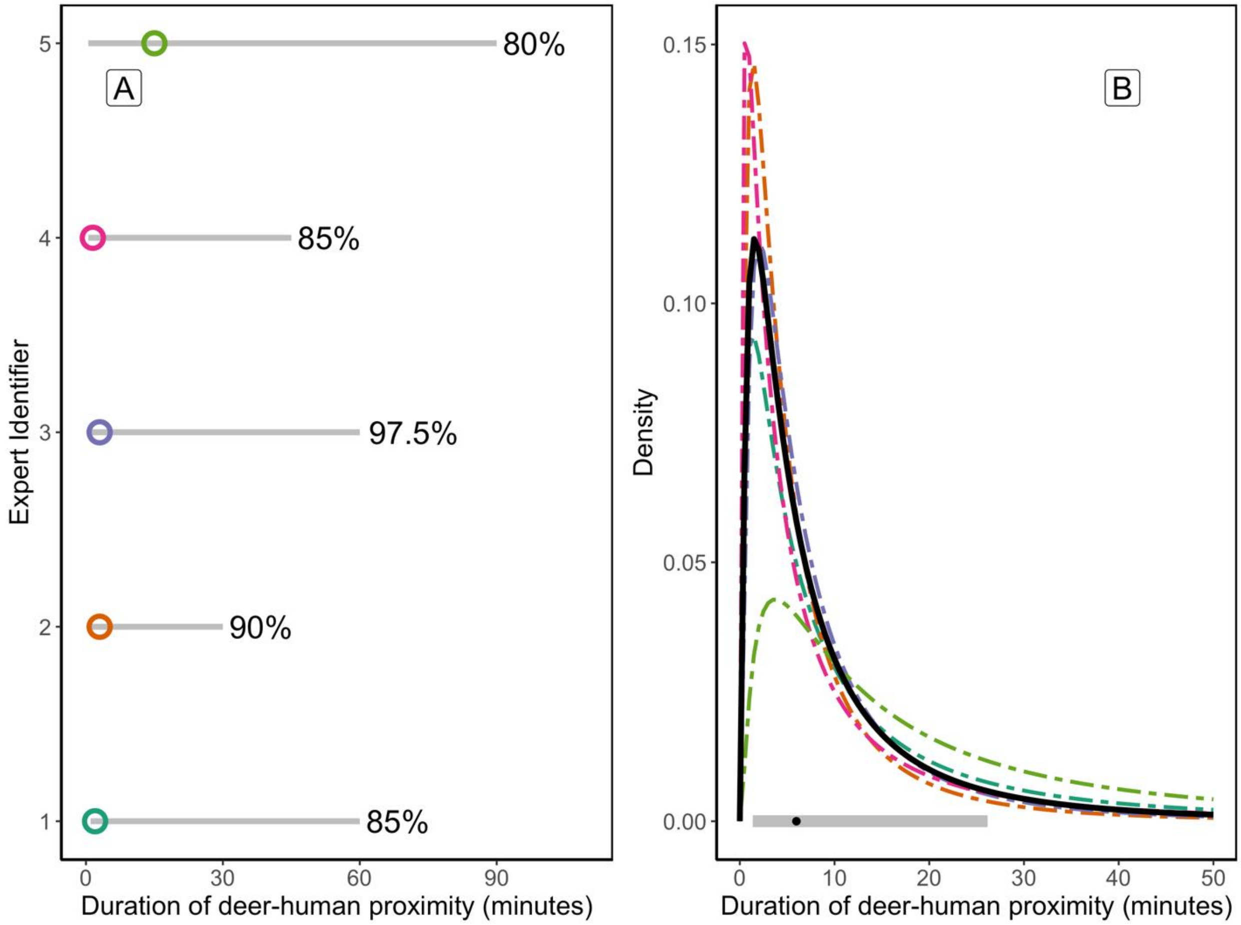

**Figure S13**: Responses by experts on the Deer Ecology panel to Question 13 to estimate the rate of a deer in proximity to another deer in an intensive captive setting (<1.5m; $\omega_{CC}$). Given these captive conditions, how many times do you expect an individual deer to be in proximity with another deer in a day on average (within 1.5m of each other)? (A) fitted log-normal probability distributions for answers from individual experts, and (B) the aggregated log-normal distribution of answers across experts. The aggregate log-normal distribution has a median proximity rate of 32.15 events per day in an intensive captive setting (80% confidence interval: 9.97–103.61 events per day; grey range along x-axis).

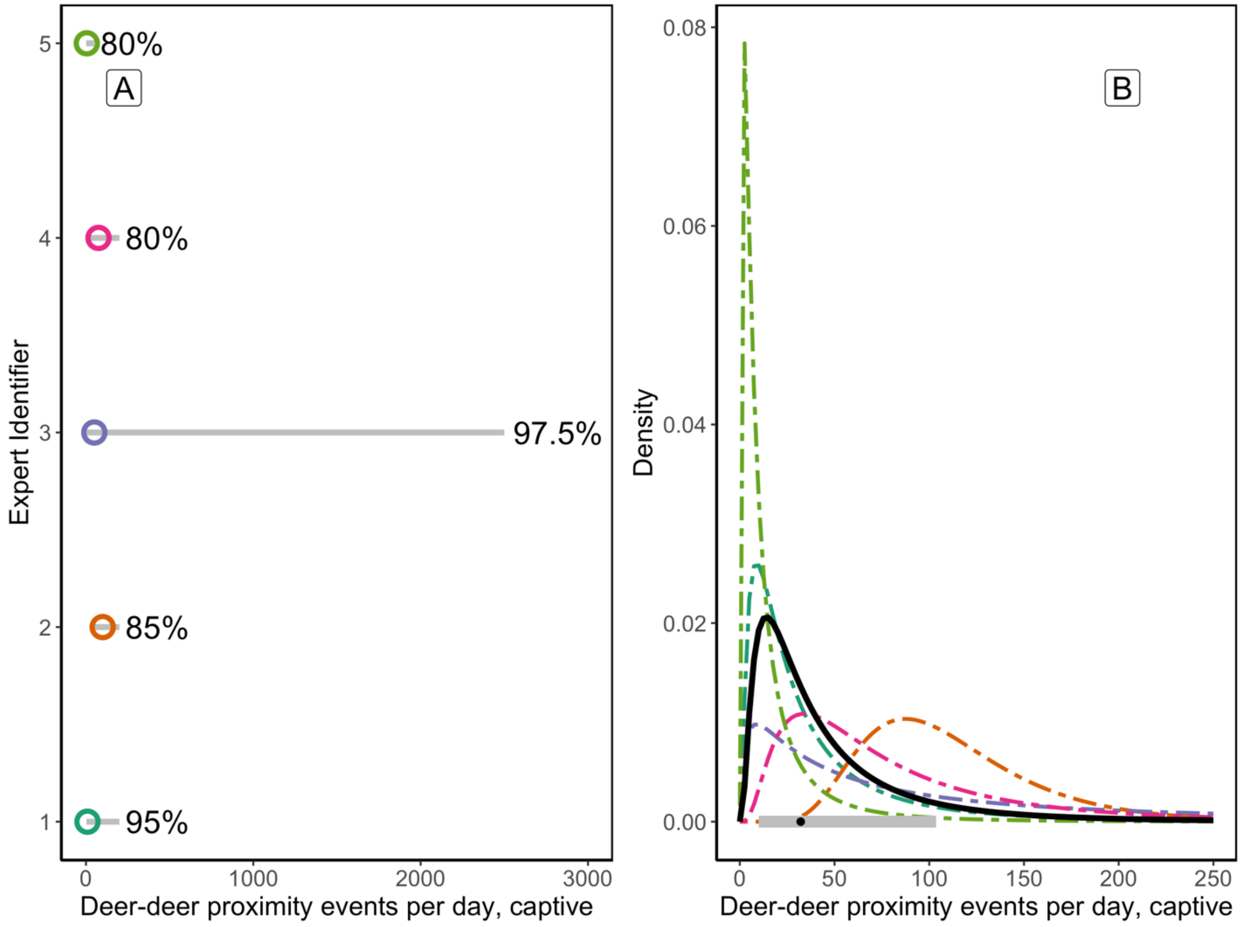
